# Resolution limit of image analysis algorithms

**DOI:** 10.1101/240531

**Authors:** Edward A. K. Cohen, Anish V. Abraham, Raimund J. Ober

## Abstract

Resolution is one of the most important properties of an imaging system, yet it remains difficult to define and apply. Rayleigh’s and Abbe’s resolution criteria^1^ were developed for observations with the human eye and had a major influence on the development of optical instruments. However, no systematic approach is available for the evaluation of the often complex image processing algorithms that have become central to the analysis of the imaging data that today is acquired by highly sensitive cameras.

Many modern imaging experiments are based on the detection of objects. Examples are localization-based superresolution experiments (PALM, STORM, etc.^2–4^), experiments to investigate the arrangement of molecular complexes on the cellular membrane such as clathrin-coated pits^5,6^, experiments tracking single particles^7,8^ or subcellular organelles^9^, etc. A specific example that we will consider in detail relates to the question of whether the distribution of clathrin-coated pits is purely random or exhibits other spatial characteristics such as clustering.

Common to the analysis of experimental data produced by such “object-based” imaging experiments is the central role that image analysis algorithms play in the identification and localization of the underlying objects, be they single molecules, clathrin-coated pits, etc. The success of such imaging experiments is, therefore, to a large extent dependent on how well these algorithms can resolve the imaged objects^10^. The assessment of such algorithms in terms of their resolution capabilities is, however, largely unexplored.

Here, we use methods of spatial statistics, which have been extensively used in different scientific disciplines^11^, to evaluate location-based image analysis algorithms. First, we show that insufficient “algorithmic resolution” can have a significant impact on the outcome of the analysis of spatial patterns which is typically carried out using the pair-correlation function or Ripley’s *K*-function^11^ (see Supplementary Material 1). For a spatial pattern that is uniformly distributed in the sense of complete spatial randomness, the pair-correlation function *g*, which describes the relationship between pairs of objects that are a distance *r* apart, is given by the identity function *g*(*r*) = 1, *r* > 0. Ripley’s *K*-function, which describes the expected number of objects within a distance *r* of an arbitrary point, is given by *K*(*r*): = *πr*^2^, *r* > 0, for a completely spatially random pattern. This implies that the related *L*(*r*) − *r* function, where 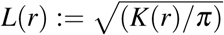 for *r* > 0, is constant and equal to zero,

The clathrin-coated pit imaging data of Figure 1a was processed using several established algorithms to determine the locations of the pits (Figure 1b,c), which were then further analyzed by plotting the estimated 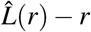 function (Figure 1d). The analysis appears to show that the pits are not distributed in a completely spatially random fashion as the 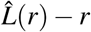 plot is not equal to zero for all the processing schemes, thereby suggesting a non-uniform arrangement of the pits on the plasma membrane. To understand this behavior, we simulated clathrin-coated pits that are located according to a completely spatially random distribution (Figure 1e). Estimating and analyzing the locations of these simulated pits (Figure 1f,g) in the same fashion as done for the experimentally acquired data reveals that the resulting 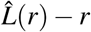 plots show remarkable similarity with those obtained from the experimentally acquired data (Figure 1h). Importantly, these plots do not show a constant value of 0 as would be expected for completely spatially random data. This suggests that the deviations from the expected constant appearance of the 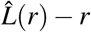 function are due to effects of the data analysis rather than being a property of the distribution of the clathrin-coated pits.

**Figure 1.**
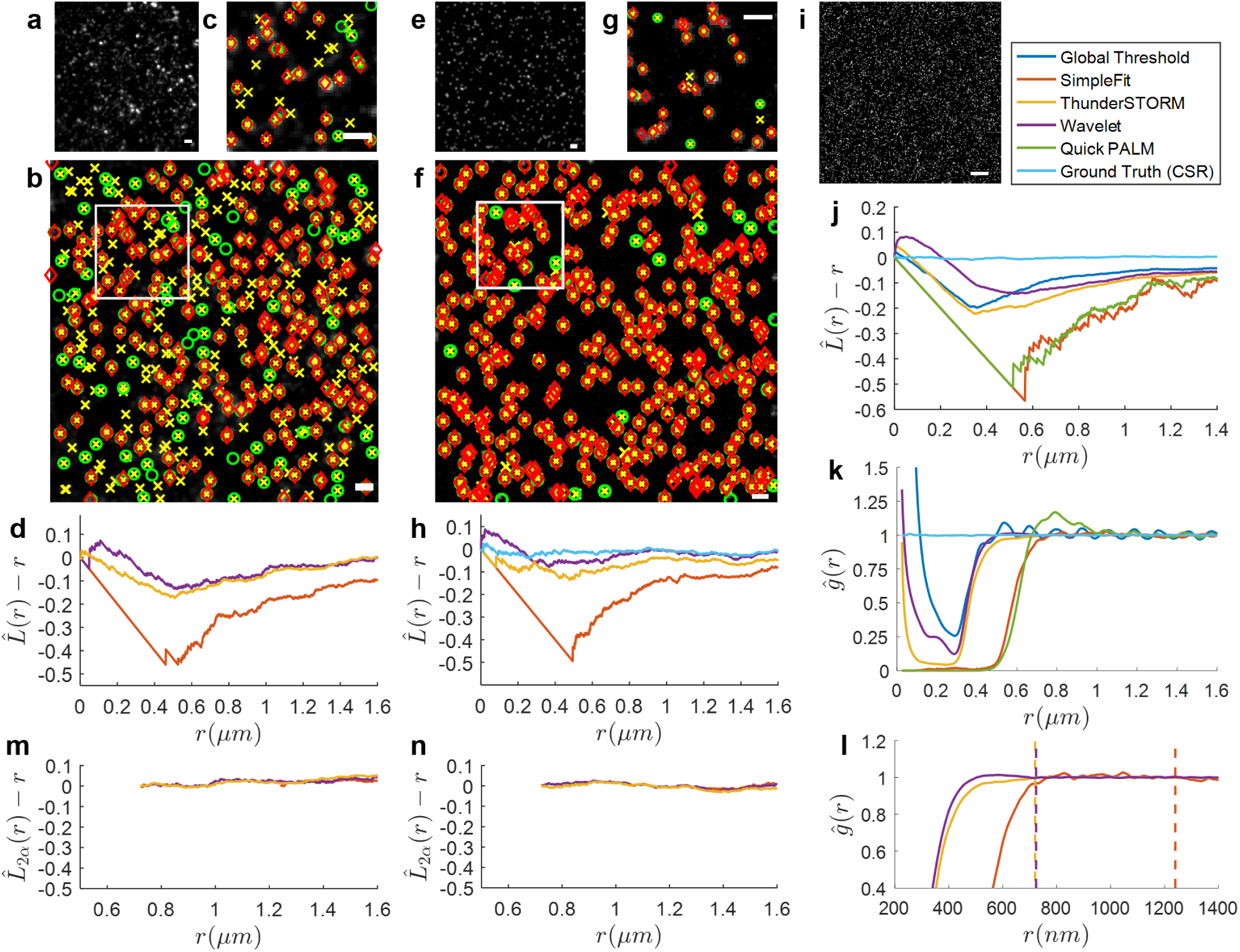
(**a**) Fluorescence microscopy image of clathrin-coated pits on the membrane of an HMEC-1 cell. Scale bar = 1*µm*.(**b**) Location estimates obtained by applying three image analysis approaches to a: fitting Gaussian profiles to clathrin-coated pits detected by wavelet-filtering (◊), using the SimpleFit software (○), and using the ThunderSTORM software (×). Scale bar = 1*µm*. (**c**) Magnified view of the region marked in **b**. Scale bar = 1*µm*. (**d**) 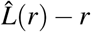 plots calculated based on the localizations shown in **b** appear to indicate that clathrin-coated pits are not distributed in a completely spatially random manner since the 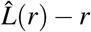 plots deviate significantly from 0 for each of the analysis approaches shown in **b**. (**e**) Simulated image of clathrin-coated pits located at completely spatially random locations. Experimental and imaging parameters similar to **a** were used for the simulation: pixel size = 6.45*µm* × 6.45*µm*, magnification = 63, and background = 100 photons/pixel. Each clathrin-coated pit was simulated using a Gaussian profile with *σ* = 120*nm* and total photon count uniformly distributed between 500 to 2000 photons. 419 clathrin-coated pits were simulated in a 200 × 200 pixel image. Scale bar = 1*µm*. (**f**) Location estimates obtained using the image analysis approaches shown in b applied to e. Scale bar = 1*µm*. (**g**) Magnified view of the marked region in **f**. Scale bar = 1*µm*. (**h**) 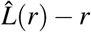 plots calculated based on the localizations shown in **f** also results in significant deviations from 0 for a completely spatially random distribution of locations. (**i**) A sample simulated image of the data set analyzed to obtain the results shown in **k** and **l**. Each image consists of 2500 molecules positioned at completely spatially random locations over a 50*µm* × 50*µm* region. The following numerical parameters were used to generate each image: pixel size = 13*µm* × 13*µm*, magnification = 100, numerical aperture = 1.3, wavelength = 525nm. Each molecule was simulated using an Airy profile with a total of 1000 photons. Scale bar = 5*µm*. (**j**) 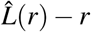 plots calculated based on localizations obtained from various image analysis approaches applied to **i** exhibit different behaviors indicating different resolving capabilities. (**k**) Pair-correlations calculated based on the localizations obtained using the image analysis approaches shown in **j** applied to a dataset containing 2000 images generated similar to **i**. These results are used to estimate the algorithm resolution limit 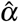 (see Supplementary Material 6, 7). The estimated algorithm resolution limits are as follows: wavelets-based algorithm 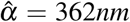, SimpleFit algorithm 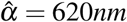, and ThunderSTORM algorithm 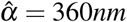. (**l**) Magnified view of the results shown in **k** with the values corresponding to 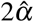 marked by dashed vertical lines. (**m**) Resolution-corrected 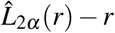 plots calculated based on the results for 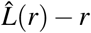 shown in **d** and corrected using the 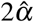 values shown in **l**. (**n**) Resolution-corrected 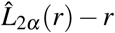 plots calculated based on the results for 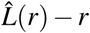 shown in **h** and corrected using the 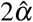 values shown in **l** no longer show significant deviations from 0 for a completely spatially random distribution of locations.

To further understand this phenomenon, we investigated whether the observed effects might be due to the different capabilities of the image processing algorithms to resolve clathrin-coated pits. To do this, we theoretically analyzed the impact of limited “algorithmic resolution” on the pair-correlation and the *L*(*r*) − *r* functions (see Supplementary Material 2). We modeled the effect of an algorithm not being able to distinguish objects that are spaced closer than a certain cut-off distance. If the objects are located in a completely spatially random fashion, the resulting *L*(*r*) − *r* function has an appearance similar to that observed in the analysis of the clathrin-coated pit data (see Supplementary Material 2.2). These observations indicate that the resolving capabilities of image analysis algorithms need to be taken into consideration when analyzing object-based imaging data.

Importantly, this analysis also suggests that the resolving capabilities of an image processing approach can be characterized by the deviation from the expected spatial analysis results for objects that are simulated with a completely spatially random location pattern. We therefore determine the *algorithmic resolution limit α* of a particular object-based image analysis algorithm by using this algorithm to estimate the locations of objects that are simulated with completely spatially random positions. Resolution effects up to a distance of *α* impact the pair-correlation function for distances up to 2*α* (see Supplementary Material 2.3). Therefore, the algorithmic resolution limit *α* is then defined as half the distance in the pair-correlation function of the estimated object locations beyond which the graph exhibits a constant plot with value 1.

We analyzed a number of well established algorithms and found that their algorithmic resolution limits can vary significantly (Figure 1l). In fact, some of these algorithms are affected by algorithmic resolution well beyond the resolution limit that is predicted by Rayleigh’s criterion, which is around 250–300 nm for the imaging conditions in Figure 1. The ThunderSTORM software^12^ has the lowest algorithmic resolution limit of 360nm, whereas the SimpleFit algorithm^13^ has an algorithmic resolution limit of 620nm, almost twice that of the ThunderSTORM software.

Our analysis has also revealed shortcomings in some established algorithms beyond the impact of algorithmic resolution. Two of the algorithms, QuickPALM^14^ and the global-threshold-based algorithm (see Methods), exhibit oscillatory behavior in the pair-correlation function even for very large distances. Upon further investigation, we found that these algorithms preferentially identify objects located towards the center of the pixels (see Supplementary Figure 1). As a result, the algorithmic resolution limit of these algorithms is taken as infinite or not defined.

Using completely spatially random data as a basis to analyze the resolution capability and to define the algorithmic resolution limit of object-based image analysis algorithms allows us to probe random configurations of object locations. Therefore, the concept of the algorithmic resolution limit also has applicability to object arrangements that are non-stochastic. As illustrated in Figure 2a (see also Supplementary Figure 2), the deterministically arranged object locations that could be reliably identified coincide with those locations that are spaced at a distance larger than the algorithmic resolution limit.

**Figure 2.**
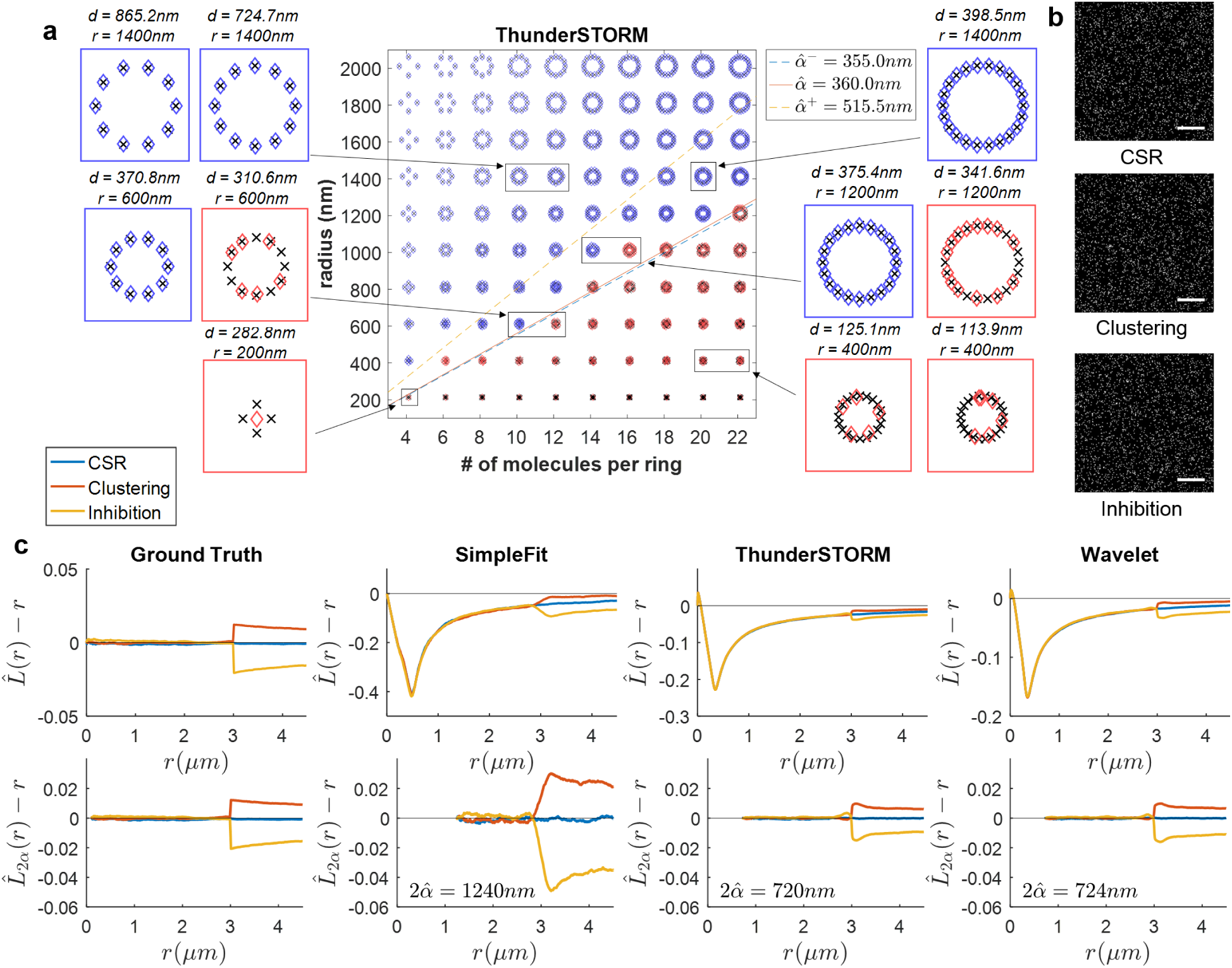
(**a**) Application of the algorithmic resolution limit to the analysis of non-stochastic data illustrated using images of deterministic structures. Each structure consists of single molecules positioned evenly around the edge of a ring (×). Localizations were obtained by analyzing the image corresponding to each structure using the ThunderSTORM software (◊). Localizations corresponding to structures where all constituent molecules were accurately identified and localized to within 10nm of the true location are shown in blue. Localizations corresponding to structures where one or more molecules were either not identified or where the localization deviated by more than 10nm from the true location are shown in red. Magnified views of some structures are shown with the radius of the corresponding ring (*r*) and the distance between adjacent molecules on the edge of the ring (*d*) indicated above each magnified view. All molecules of structures where the spacing between adjacent molecules is greater than the algorithmic resolution limit of ThunderSTORM are accurately identified and localized to within 10nm of the true location. The solid line corresponding to 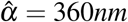 indicates the algorithmic resolution limit for ThunderSTORM. The dashed lines on either side of the solid line indicate the bootstrapped 80% confidence interval for the estimate of *α* (see Supplementary Material 6.2). Results obtained by analyzing the same images using other approaches are provided in Supplementary Figure 2. (**b**) Sample images from three data sets that were analyzed to obtain the results shown in **c**. The three data sets were generated with the following spatial distribution of molecules (see Methods): completely spatially random (CSR) distribution, random distribution with a preferred spacing ranging from 2990nm to 3010nm between molecules (clustering), and random distribution with molecules avoiding spacings between 2990nm to 3010nm of each other (inhibition). Scale bar = 10*µm*. (**c**) 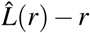 plot compared to the corresponding resolution-corrected 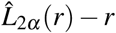 plot calculated based on localizations obtained by analyzing the three data sets illustrated in **b**. For each analysis approach, the value corresponding to 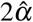 indicated in **Fig. 1l** is used to calculate 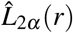. Results show that deviations from 0 in the 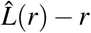 plot are corrected for completely spatially random distributions of locations when the algorithmic resolution limit is taken into account. Results for distributions with clustering or inhibition spacings between molecules still show corresponding deviations from 0 in the corrected 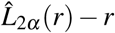 results.

The question immediately arises, how the algorithmic resolution limit of an image analysis algorithm impacts the analysis of experimental data. For example, it is important to quantitate how many objects remain unaffected by resolution effects when the imaging data is analyzed using an algorithm with algorithmic resolution limit *α*. As shown in Supplementary Material 3, the probability that an object is unaffected by resolution effects is given by 1 − *G*_0_(*α*), where *G*_0_ is the nearest-neighbor probability distribution function which describes the probability that an arbitrary object is at a distance less than *α* from its nearest neighboring object (see Supplementary Material 1). If the objects are located according to a completely spatially random distribution, *G*_0_(*α*) = 1 − exp(−*λ*_0_*πα*^2^), where *λ*_0_ is the density of the object locations. Therefore, in a highly dense object pattern, not suprisingly, the probability that an object is not affected by algorithmic resolution is severely reduced.

For example, consider cellular membrane receptor clusters distributed in a completely spatially random fashion with a density of 1 cluster per square micrometer (as in Figure 1e). When analyzing the location of such protein clusters using an algorithm with algorithmic resolution limit *α* = 360*nm* (e.g., the ThunderSTORM software^12^), the probability of the location being unaffected by resolution is 66.5%. However, when analyzing the cluster locations using an algorithm with *α* = 620*nm* (e.g., the SimpleFit algorithm^13^), the probability of the location being unaffected by resolution is 29.9%. Thus, the difference in algorithmic resolution between the two algorithms can have drastic effects on the analysis of the data. Further, it is only for cluster densities of 0.1 clusters per square micrometer that the probability of a cluster being unaffected by resolution will be above 95% (with 96.0%) for the algorithm with *α* = 360*nm*. However, this probability decreases significantly to 88.6% for the algorithm with *α* = 620*nm*.

Localization-based superresolution methods use repeat stochastic excitation of small subsets of the fluorophores in a sample^4^. The question therefore arises how small these subsets need to be in order for a large fraction of the single molecules/objects to not be affected by the algorithmic resolution limit of the analysis step. For a tubulin dataset^10^ (also see Supplementary Materials 5.4) the probability that an object is unaffected by resolution is 86.4% when analyzed using an algorithm with algorithmic resolution limit of *α* = 360*nm* (e.g., ThunderSTORM) for *q* = 1/2014, where *q* is the probability of an object appearing in any given frame of the dataset. However, changing *q* to 1/10000 increases the probability to 96.8% whereas for *q* = 1/1000 the probability that a single molecule is unaffected by resolution is decreased to 71.2%. Interestingly, in order to achieve the probability of 95% in a classical single molecule experiment where all single molecules are activated and imaged in a single acquisition, an algorithmic resolution limit of 0.8*nm* would be required, which is well beyond what is currently achievable, thus illustrating the power of using stochastic excitation for localization-based single molecule superresolution experiments.

We have seen that algorithmic resolution can significantly distort Ripley’s *K*-function. However, knowing the algorithmic resolution limit *α* of an algorithm allows us to define a resolution-corrected Ripley’s *K*_2_*_α_*-function and resolution-corrected *L*_2_*_α_*(*r*) − *r* for *r* ≥ 2*α* (see Supplementary Material 8). Figure 2c shows that inhibition and clustering can be correctly identified with the resolution-corrected *L*_2_*_α_*(*r*) − *r* function if they occur at distances above 2*α* for object-based imaging data analyzed with an algorithm of resolution limit *α*.

If the clathrin-coated pit data of Figure 1a is analyzed using an algorithm with algorithmic resolution limit *α* = 360*nm* (e.g., the ThunderSTORM software) and the estimated locations processed with the resolution-corrected *L*_2_*_α_*(*r*)−*r* function, the data shows that there is no significant deviation of the clustering behavior from complete spatial randomness beyond the distance of 2*α* = 720*nm* (Figure 1m, n). This indicates that at distances above twice the algorithmic resolution limit for the individual algorithms the clathrin-coated pit locations do not show any deviation from complete spatial randomness.

Resolution has been analyzed in microscopy going back to the classical criteria by Rayleigh and Abbe. Those criteria address the performance of the imaging optics. Using an information theoretic approach, Rayleigh’s resolution criterion was generalized and put in the context of modern imaging where data consists of noise-corrupted photon count measurements acquired through quantum-limited detectors^15^. A resolution measure based on the Fourier ring coefficient was introduced that can be computed directly from an acquired image and takes into account the standard deviation with which a single molecule can be localized^16^. Common to these recent approaches is that they assume that the image analysis algorithms do not have resolution limitations themselves. Here we have introduced a methodology to systematically assess the algorithmic resolution limit of object-based image analysis algorithms and to evaluate the impact of the limitations on the analysis of microscopy data. We hope that the approaches presented here will contribute to a systematic evaluation of such algorithms that are of relevance not only to microscopy applications but to other object-based imaging scenarios such as those arising, for example, in astronomy.

## Additional information

### Acknowledgements

The authors would like to thank Professor Niall Adams, Department of Mathematics, Imperial College London for valuable discussions and input on implementing change point methods to estimate algorithmic resolution limits, and Sreevidhya Ramakrishnan, Department of Biomedical Engineering, Texas A&M University for preparing the cellular samples and acquiring the experimental data of clathrin-coated pits. This work was supported in part by the National Institutes of Health (R01GM085575).

### Author Contributions

E.A.K.C and R.J.O conceived the methodology presented here, developed the mathematical theory underlying the methodology, and wrote the manuscript. A.V.A performed the calculations and prepared the figures for the results.

### Correspondence

Please direct all correspondence to or

### Data availability

The data is available on request.

### Code availability

The software related to the analysis presented in this paper is available at https://github.com/eakcohen/algorithmic-resolution and http://www.wardoberlab.com/software^1^.

## Methods

### Preparing HMEC-1 cells for fluorescence imaging

HMEC-1 cells were fixed using 1.7% (w/v) Paraformaldehyde (Electron Microscopy Sciences) at room temperature and permeabilized by incubation with 0.02% (w/v) saponin in phosphate buffered saline (PBS) for 10 minutes at room temperature. Cells were then pre-blocked with 3% BSA in PBS, incubated with anti-Clathrin primary antibody (mouse monoclonal X22, diluted 1000-fold in 1% BSA/PBS, Abcam) for 25 minutes at room temperature, and treated with goat serum diluted 50-fold. Bound primary antibody was detected by treatment with Alexa 555-labeled anti-mouse IgG (diluted 750-fold in 1% BSA/PBS, Invitrogen) for 25 minutes at room temperature. Cells were washed twice with PBS between each incubation and finally immersed in 1.5 mL of 1% BSA/PBS prior to imaging.

### Fluorescence microscopy imaging

Fixed HMEC-1 cells were imaged with a Zeiss (Axiovert 200M) inverted epifluorescence microscope fitted with a 63x (1.4 NA) Plan Apo objective (Carl Zeiss) using a CCD camera (Orca ER, Hamamatsu). The sample was illuminated using a broadband LED illumination (X-Cite 110LED, Excelitas Technologies) filtered through a standard Cy3 filterset (Cy3-4040C-ZHE M327122 Brightline, Semrock). Signal from the sample was also filtered through this filterset before being acquired by the camera.

### Generating location distributions

#### Completely spatially random distribution of locations

For a completely spatially random distribution of locations in an *Sµm* × *Sµm* region, the (*x*,*y*) coordinate for each location was obtained by drawing realizations of independent random variables *X* and *Y*, each uniformly distributed with probability density functions *p_X_* (*x*) = *p_Y_* (*y*) = 1/*S*, 0 ≤ *x*, *y* ≤ *S*.

#### Locations for deterministic structures

Deterministic structures consist of *D* molecules located at evenly spaced points on the circumference of a circle of radius *r*. The location of the *d^th^* point 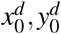 is given by 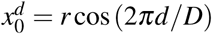 and 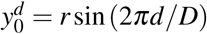, where *d* = 1,…,*D*.

#### Random distribution of locations exhibiting a preferred spacing (clustering)

A set of *N* random locations, denoted by Δ, such that *N_c_* pairs of those locations are spaced at distances between *r*_min_ and *r*_max_ is generated by combining three subsets of locations so that 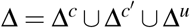. The subset 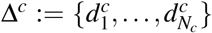 consists of completely spatially random locations 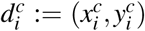. The subset 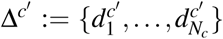 consists of locations corresponding to Δ*^c^*, where each location 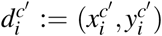, is calculated as

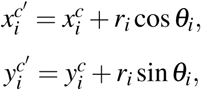

The distance *r_i_* between 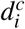 and 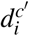 is uniformly distributed between *r*_min_ and *r*_max_, and θ*_i_* is uniformly distributed between 0 and 2*π*. The subset Δ*^u^* is an additional completely spatially random distribution of *N_u_* = *N* − 2*N_c_* locations. For the simulated images analyzed to obtain the results in Figure 2, the following values were used: *N* = 2500, *N_c_* = 250, *r*_min_ = 2990*nm*, and *r*_max_ = 3010*nm*.

#### Random distribution of locations avoiding specific spacings (inhibition)

A set of locations, Δ := {*d*_1_,…,*d_N_*}, in which no two locations are spaced between *r*_min_ and *r*_max_ of each other is generated as follows. For *i* = 1,…,*N*, the *i^th^* location *d_i_* is drawn from a completely spatially random distribution. The *i^th^* point is not added as a location if *r*_min_ ≤ *d_i j_* ≤ *r*_max_ for some 1 ≤ *j* ≤ *i*, where *d_i j_* denotes the distance between the *i^th^* and *j^th^* locations. For the simulated images analyzed to obtain the results in Figure 2, the following values were used: *N* = 2500, *r*_min_ = 2990*nm* and *r*_max_ = 3010*nm*.

### Simulating Images

#### Simulating an image of clathrin-coated pits

When simulating an image of clathrin-coated pits, the detector is modeled as a set of pixels {*C*_1_,…,*C_K_*}. The photon count detected in the *k^th^* pixel is modeled as 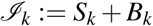, where *S_k_* and *B_k_* are both Poisson random variables^17^. The total photons detected at the *k^th^* pixel from all clathrin-coated pits within the region represented by the image is denoted by *S_k_*. The background photon count *B_k_* has a mean of *B* = 100 photons/pixel.

The mean of *S_k_* is given by

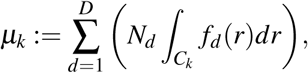
where *D* is the total number of clathrin-coated pits in the image, *N_d_* denotes the total number of photons detected from the *d^th^* clathrin-coated pit, *C_k_* denotes the area of the *k^th^* pixel, and *f_d_* denotes the photon distribution profile for the *d^th^* clathrin-coated pit. For the simulated image of clathrin-coated pits in Figure 1, *D* = 419 and *N_d_* had values uniformly distributed between 500 to 2000 photons.

For each clathrin-coated pit, *f_d_* is modeled as a Gaussian profile given by

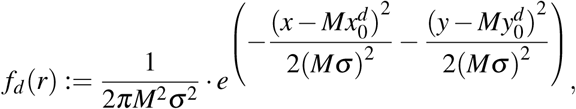
where, *M* denotes magnification, *σ* denotes the width of the Gaussian profile, and 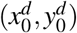 denotes the center of the *d^th^* clathrin-coated pit. The coordinate 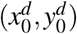 is drawn from a completely spatially randomly distributed set of *D* locations generated as described above.

#### Simulating images of single molecules

When simulating an image of single molecules, the detector is again modeled as a set of pixels {*C*_1_,…,*C_k_*}. The photon count detected at the *k^th^* pixel is modeled as a Poisson random variable with mean given by,

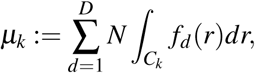
where *N* denotes the total number of photons detected from the molecule, *C_k_* denotes the area of the *k^th^* pixel, and *f_d_* denotes the photon distribution profile for the *d^th^* molecule. For each molecule, *f_d_* is modeled as an Airy profile given by

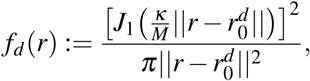
where *J*_1_ denotes the first-order Bessel function of the first kind, 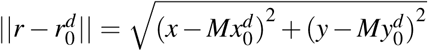, 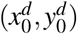 denotes the location of the *d^th^* molecule, and M denotes the magnification of the optical system. *κ* is calculated as *κ* = 2*πN_a_*/*λ*, where *N_a_* denotes the numerical aperture, and *λ* denotes the wavelength of the photons emitted by the molecule. The following values were used for simulating all images of single molecules: *N_a_* = 1.3, *λ* = 525*nm*, *M* = 100.

### Image Analysis

#### List of image anlaysis approaches

The following is a list of the image analysis approaches that were used:

- **Wavelet:** Detects molecules or clathrin-coated pits using wavelet-filtering^18^ and estimates their locations by fitting Airy profiles to the detected molecules or Gaussian profiles to the detected pits. Further details are provided below.
- **Global Thresholding:** Detects molecules or clathrin-coated pits by identifying pixels above a threshold value and estimates their locations by fitting Airy profiles to the detected molecules or Gaussian profiles to the detected pits. Further details are provided below.
- **SimpleFit:** Detects and localizes molecules using the default settings of the software package available from^19^.
- **ThunderSTORM:** Detects and localizes molecules using the default settings of the software package described in^20^.
- **QuickPALM:** Detects and localizes molecules using the default settings of the software package described in^21^.

#### Identifying single molecules or clathrin-coated pits by wavelet-filtering

The image was filtered using the product of two consecutive wavelet transforms as described in^18^. Each isolated set of one or more edge-connected pixels obtained from the filtering was identified as a region of the image containing an individual molecule or clathrin-coated pit. For the subsequent localization of that molecule or clathrin-coated pit, a 5 × 5 pixel region centered on the average pixel coordinate of the corresponding set of identified pixels was used.

#### Identifying single molecules or clathrin-coated pits by global-thresholding

The image was thresholded using 25% of the maximum pixel intensity in the dataset as the threshold value. Each isolated set of one or more edge-connected pixels obtained from the thresholding was identified as a region of the image containing an individual molecule or clathrin-coated pit. For the subsequent localization of that molecule or clathrin-coated pit, a 5 × 5 pixel region centered on the average pixel coordinate of the corresponding set of identified pixels was used.

#### Localizing clathrin-coated pits and single molecules identified by wavelet-filtering or global-thresholding

Each clathrin-coated pit and single molecule was localized by fitting a Gaussian and Airy profile, respectively, to the corresponding 5 × 5 pixel image identified using either wavelet-filtering or global-thresholding as described above. An initial location estimate and an initial value for the *σ* parameter denoting the width of a Gaussian profile was calculated for each clathrin-coated pit or single molecule by applying the approach described in^22^ to the corresponding image. An initial value for the *κ* parameter of the Airy profile was calculated as 1.323/*σ*. The background associated with each clathrin-coated pit or single molecule was taken as the median of the intensities in the edge pixels of the corresponding 5 × 5 pixel image. An initial estimate of the photon count detected from each molecule or clathrin-coated pit was taken as the sum of the pixel intensities in the corresponding image after subtracting the background.

Airy or Gaussian profiles with initial values for the various parameters calculated as described above were fitted to each 5 × 5 pixel image using a least-squares estimator to obtain the final location estimates. The location parameters (*x*_0_,*y*_0_), width parameter (*σ* when fitting Gaussian profiles and *κ* when fitting Airy profiles), and the total photon count were estimated for each clathrin-coated pit or single molecule.

#### Estimating *L*(*r*) − *r* using localizations from one image

When estimating *L*(*r*) − *r* for a set of localizations obtained by analyzing an image of either clathrin-coated pits or single molecules, 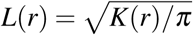 for *r* > 0. *K*(*r*) denotes the Ripley’s *K*-function, defined as

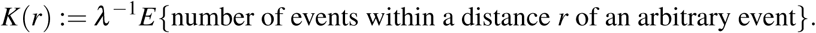

The estimator for *K*(*r*) is given by

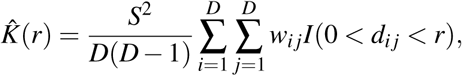
where *S*^2^ denotes the area in the object space corresponding to the image being analyzed, *w_i j_* denotes the Ripley’s isotropic edge correction weights^23^, *D* denotes the total number of localizations, and *d_i j_* denotes the distance between the *i^th^* and *j^th^* localizations. The indicator function is defined as

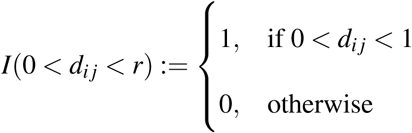

Estimating *L*(*r*) − *r* using localizations from multiple images

When estimating *L*(*r*) − *r* for a particular image analysis approach from a total of *D* localizations distributed among *B* images,

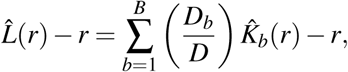
where *D_b_* denotes the number of localizations obtained from the *b^th^* image and 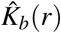 denotes estimates of the Ripley’s K-function calculated using the localization obtained from that image.

#### Estimating pair-correlations for an image analysis approach

Estimates of the pair-correlation results for an image analysis approach, denoted as *a*, were calculated by a weighted averaging of pair-correlation estimates from multiple simulated images as follows. A total of *B* images containing *D* single molecules were simulated. A set of localizations of size 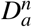 were obtained by applying analysis approach *a* to the *b^th^* image, for *b* = 1,…,*B*. Pair-correlations estimates 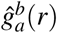 were calculated independently for each set of 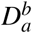 localizations using a MATLAB implementation of the approach in^24^. The weighted-average pair-correlation estimates for each analysis approach *a* is then calculated as

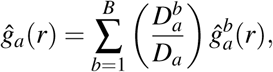
where 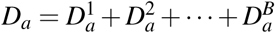. The pair-correlation results shown in Figure 1 were calculated using B = 2000 images containing D = 250,000 molecules.

#### Determining *α* from pair-correlations for an image analysis approach

The radius of correlation for analysis approach *a* is defined as

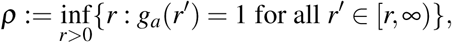
when *a* analyzes completely spatially random data. To estimate *ρ* from 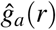, the scheme presented in Supplementary Material Section 6 was implemented. For a set 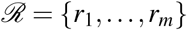 of finely, regularly spaced proposal values of *ρ*,

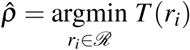
where

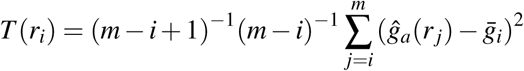
and 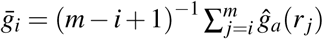. Algorithmic resolution limit *α* is then determined as 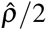.

#### Calculating *L*_2_*_α_* (*r*) − *r* for an image analysis approach

For the algorithmic resolution limit *α* for a specific image analysis approach determined as described above, *L*_2_*_α_* (*r*) − *r* is calculated as

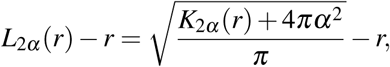
where *K*_2_*_α_* (*r*) = *K*(*r*) − *K*(2*α*). See Supplementary Material 6 for details regarding the determination of 2*α* for each image analysis approach from the corresponding pair-correlation results.

### Software

ROI identification using wavelet-filtering or global-thresholding followed by fitting with either Airy or Gaussian profiles was performed using custom programs developed with the MIATool software framework^25^ in Java. The ThunderSTORM^20^ and SimpleFit^19^ software packages were used with default settings for the various options within the software. The QuickPALM^21^ software was used with the FWHM=2 setting to match the width of the single molecule or clathrin-coated pit being localized. Calculations for *L*(*r*) − *r*, pair-correlations, and *L*_2_*_α_* (*r*) − *r* were performed using custom-developed scripts in the MATLAB programming environment (The MathWorks, Inc., Natick, MA). All figures were similarly prepared using MATLAB.

## Footnotes

1 The software will be publicly released on acceptance of this paper

